# Assessing Immune Stability in Space: Haematological and T Cell Profiling of Astronauts During the MESSAGE Science Mission in Axiom-3 on the ISS

**DOI:** 10.1101/2025.04.09.647940

**Authors:** Cihan Tastan, Gamze Gulden, Ozge Demir, Ebru Cam, Fatmanur Erkek, Berranur Sert, Tayfun Gozler, Ipek Yuksel, Busra Tekirdagli

## Abstract

Microgravity presents unique challenges to human physiology, particularly the immune system, during spaceflight. T lymphocytes, key components of adaptive immunity, play a vital role in immune regulation, yet the effects of microgravity on T-cell function and gene expression remain incompletely understood. This study, conducted as part of the MESSAGE (Microgravity Associated Genetics) Science Mission in the Axiom-3 mission (https://www.nasa.gov/mission/station/research-explorer/investigation/?#id=9100). We aimed to investigate how microgravity influences immune cell responses by analysing blood samples from three astronauts between pre-flight and during their space mission on the ISS (post-launch Day 4, Day 7, and Day 10 in ISS). Hemogram analysis revealed no statistically significant differences in leukocyte, erythrocyte, haemoglobin, and haematocrit levels between pre-flight and in-flight samples, suggesting a stable haematological profile under microgravity conditions. T cell subpopulation analysis indicated fluctuations in effector memory T cells (TemEARLY and TemLATE), but these changes did not reach statistical significance. These findings suggest that, within the short duration of spaceflight, haematological and immune parameters remain largely stable, underscoring the need for further research into long-term immune adaptations in microgravity. As a next step, transcriptomic analysis will be performed to identify microgravity-associated gene candidates, and CRISPR-based knock-out of the genes in T cells will be generated to explore their functional roles in microgravity-induced immune modulation.

## 1. Introduction

Gravity is a fundamental force that significantly influences the structure and function of living organisms on Earth. It affects cellular behaviour, tissue organization, and overall physiology. Understanding how the absence of gravity, or microgravity, affects human biology is essential for the advancement of long-duration space missions. Microgravity has been shown to influence various physiological systems, including the cardiovascular, musculoskeletal, and immune systems [1]. Of particular interest is the impact of microgravity on the immune system, as immune dysregulation during spaceflight poses a significant risk to astronaut health.

The immune system is composed of a complex network of cells and molecules that protect the body from infections and other harmful agents. Among these cells, T lymphocytes play a critical role in adaptive immunity. T cells are responsible for destroying infected host cells, producing cytokines, activating other immune cells, and regulating immune responses [2]. Numerous studies have demonstrated that spaceflight can alter immune cell distribution, reduce T cell proliferation, and impair cytokine production, leading to a compromised immune response [3,4]. One of the key challenges faced by astronauts is immune dysregulation caused by the unique stressors of spaceflight. These stressors include microgravity, increased radiation exposure, psychological stress, and altered circadian rhythms [5]. Research aboard the International Space Station (ISS) has provided valuable insights into how microgravity affects immune function. Studies have shown that leukocyte distribution, cytoskeletal structure, and the functional capacity of immune cells are disrupted in microgravity, resulting in decreased interleukin-2 (IL-2) production and increased tumour necrosis factor-alpha (TNF-α) production [6]. Moreover, microgravity has been associated with increased reactivation of latent viruses, further highlighting the vulnerability of the immune system during space missions [7]. Despite significant advancements in our understanding of how spaceflight affects the immune system, the molecular mechanisms underlying these changes remain poorly understood. There is a growing interest in identifying gravity-sensitive genes and understanding how changes in gene expression may contribute to immune dysregulation. Recent advances in transcriptomics and gene editing technologies, such as CRISPR, provide new opportunities to explore the genetic basis of immune changes in microgravity [8].

In this study, we aimed to investigate the effects of microgravity on the immune system by analysing haematological and T cell subpopulation changes in astronauts participating in the Axiom-3 mission as part of the MESSAGE (Microgravity Associated Genetics) Science Mission. Blood samples were collected at multiple time points (post-launch Day 4, Day 7, and Day 10 on the ISS) to assess immune alterations in response to spaceflight. By conducting comprehensive hemogram and flow cytometry analyses, we sought to determine whether microgravity induces significant changes in immune cell populations. The results indicated no statistically significant alterations in haematological parameters or T cell subpopulations, suggesting that short-term exposure to microgravity does not lead to major immune system dysregulation. These findings highlight the stability of immune responses during short-duration space missions while emphasizing the need for further investigations into long-term immune adaptations in microgravity. Understanding these responses is critical for safeguarding astronaut health and advancing space medicine.

## 2. Material and methods

### Ethics Approval and Consent to Participate

Microgravity Associated Genetics (MESSAGE) Science Mission proposal (Study eIRB Number: STUDY00000620) has been reviewed and approved (Date of Approval: 21Sep2023) by the NASA IRB in accordance with ethical standards and the requirements of the Code of Federal Regulations on the Protection of Human Subjects (NASA 14CFR1230 and if applicable, FDA 21CFR50 and 56). MESSAGE Science Mission proposal (No: 61351342/020-275) has been also approved (Date of Approval: 03July2023) by Üsküdar University Ethical Commission.

### Isolation of Peripheral Blood Mononuclear Cells (PBMC)

Peripheral blood mononuclear cells (PBMC) were isolated from healthy adult blood samples obtained at the Üsküdar University Transgenic Cell Technologies and Epigenetic Application and Research Center (TRGENMER). Blood samples of 3cc were obtained from three healthy human donors and diluted in a 1:1 ratio with PBS (without Mg/Ca). The blood sample diluted in a 1:2 ratio with Ficoll (Paque PREMIUM 1.073, 17-5446-52) was combined. PBMCs were isolated using density gradient centrifugation. The sample combined with Ficoll was centrifuged at 800 g for 25 minutes with an acceleration of 9 and brake at 0. The Buffy coat formed after centrifugation was collected into a new tube. PBS (without Mg/Ca) was added in a 1:3 ratio (Buffy coat) and centrifuged at 400 g for 10 minutes. This process was repeated until the supernatant became clear. After centrifugation, the pellet was dissolved in 5 ml of PBS (without Mg/Ca) and centrifuged at 400 g for 5 minutes. The pellet was dissolved in RPMI Medium (CAPRICORN SCI, RPMI-A) (10% FBS + 1% penicillin/streptomycin). The 3cc blood sample was analysed for hemogram in NPISTANBUL Brain Hospital, Türkiye.

### Analysis of T Cell Sub-Populations (TN-TCM-TEM-TEF) and Immunodeficiency Profile

T cell sub-populations and Tcm-Tscm profiling studies on days 4, 7, and 10 were analysed using CD3-PC7 (Beckman Coulter, 6607100), CD4-APC-A700 (Beckman Coulter, B10824), CD8-PC5.5 (Beckman Coulter, B21205), CD45RA-ECD (Beckman Coulter, IM2711U), CD45RO-PE (Miltenyi Biotech,130-113-559), CD27-APC-A750 (Beckman Coulter, B12701) and CD62L-APC (Miltenyi Biotech, 130-113-617) antibodies. Exhaustion profiling studies on days 14 and 21 were analysed using CD3-PC7 (Beckman Coulter, 6607100), CD4-APC-A700 (Beckman Coulter, B10824), CD8-PC5.5 (Beckman Coulter, B21205), CD279 (PD1)-PE (Miltenyi Biotech, 130-117-384), CD366 (TIM3) - APC (Invitrogen, 17-3109-42) and CD223 (LAG-3) - APC-eFluor 780 (Invitrogen, 47-2239-42) antibodies. 100 μl samples were taken by pipetting and added to tubes. 1 ml of Staining Buffer was added onto the samples (Staining Buffer, 1:20-BSA:Rinsing Solution). After adding buffer, it was centrifuged at 400g for 5 minutes. After centrifugation, the supernatant was discarded, and the final volume was adjusted to 20 μl. Antibodies for the Tcm-Tscm and Exhaustion panels were added according to the manufacturer’s instructions and vortexed in the dark. After vortexing, tubes were incubated at +4^0^C for 30 minutes. After incubation, 1 ml of Staining Buffer was added to the samples in the dark. It was centrifuged at 400g for 5 minutes. By discarding the supernatant, the pellet was dissolved in 100 μl Staining Buffer and analysed by Flow Cytometry. Survival analysis was performed to control the cell viability of PBMC. Trypan blue (Biologica Industries, #03-102-1B) was applied to identify and count surviving cells. Cell counting and viability analysis were performed with the BIO-RAD TC20 Automated Cell Counter.

### Statistics Analysis

Two-tailed T-tests and Wilcoxon were performed using SPSS software or bar graphs. Outliers were not excluded in any of the statistical tests and each data point represents an independent measurement. Bar plots report the mean and standard deviation or the standard deviation of the mean. The correlation analyses were conducted using GraphPad Prism software with XY analyses, and Pearson’s correlation test was applied to assess the linear relationship between two continuous variables. The obtained p-values determined the statistical significance of these relationships, with p < 0.05 indicating a significant correlation.

## 3. Results

### Hemogram analysis

To understand the effects of microgravity on cellular aging and the immune system, hemogram analysis, and immune profiling experiments were conducted as part of the ground-based experiments. Blood samples were collected from the astronaut at multiple time points: Day 4, Day 7, and Day 10 during their stay in space. Cell counts were performed, and the samples were cryogenically preserved for further analysis. The hemogram analysis, a laboratory test evaluating the components of blood, particularly erythrocytes (red blood cells), leukocytes (white blood cells), and platelets, was carried out to assess changes in the astronaut’s health during spaceflight. These results are detailed in (as shown in Table 1 and see Fig.1). The hemogram results provide valuable insights into how space conditions affected the astronaut’s haematological profile. The counts of various blood cells at each time point (Day 4, Day 7, and Day 10) show key changes that are indicative of the body’s response to microgravity. This data is crucial for understanding how space environments influence overall immune function and cellular aging. The hemogram analysis aimed to assess the astronaut’s blood profile during spaceflight at three key time points: Day 4, Day 7, and Day 10. Leukocytes were noticeable increase from Day 4 (Mean 1.23 ± 2.05) to Day 10 (Mean 2.80 ± 0.62). While the difference between Day 4 and Day 7 was not statistically significant (p=0.109), a higher leukocyte count on Day 10 suggests immune activation. The erythrocyte count remained relatively stable between Day 4 (Mean 4.75 ± 0.75) and Day 7 (Mean 4.77 ± 0.77) but dropped slightly on Day 10 (Mean 4.23 ± 0.50). No significant differences were noted between the time points (p=0.593 and p=0.109). The haemoglobin levels remained consistent across the three time points, with no statistically significant changes (p-values >0.1). Also, there was a non-significant reduction in haematocrit on Day 10 compared to Day 4 and Day 7, potentially indicating a change in blood volume or red cell mass during spaceflight. The MCV increased slightly from Day 4 to Day 7 but then dropped on Day 10. However, these changes were not statistically significant (p=0.285). MCH and MCHC values exhibited minor fluctuations across the three time points, but no significant changes were observed. Platelet levels increased from Day 4 to Day 7 but then slightly decreased on Day 10, with no statistically significant differences (p=0.109, p=0.593). RDW parameters, which measure red blood cell distribution width, showed slight variations across the time points, but the differences were not statistically significant. HbA1c levels, a marker of long-term glucose control, remained stable across all time points, indicating no significant changes in glucose metabolism during the space mission. The data suggest that while there were slight fluctuations in some haematological parameters during spaceflight, particularly in leukocyte count and haematocrit levels, most other blood components remained stable. This highlights a relatively controlled response to the space environment in terms of blood cell production and turnover.

**Table 1:**
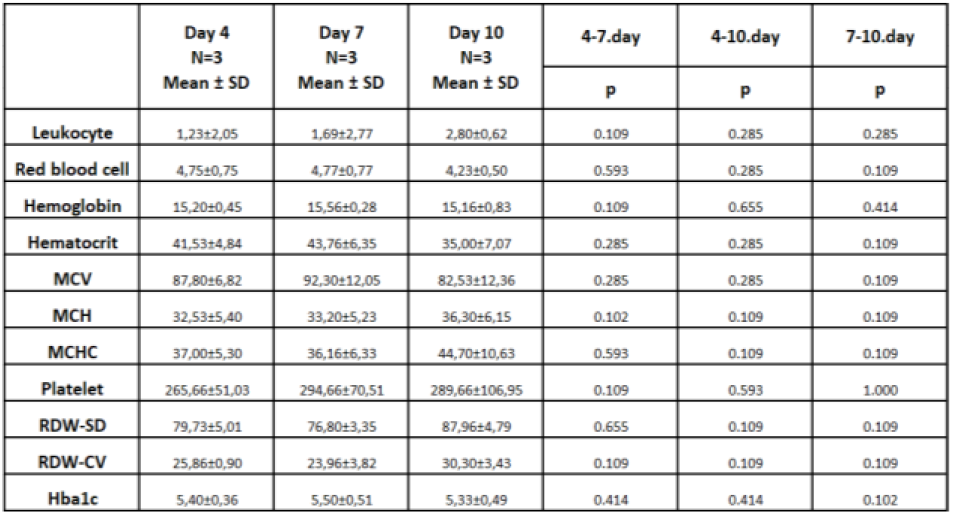
Hemogram Analysis Results (t-test)

| | Day 4<br>N=3<br>Mean ± SD | Day 7<br>N=3<br>Mean ± SD | Day 10<br>N=3<br>Mean ± SD | 4-7.day | 4-10.day | 7-10.day |
| --- | --- | --- | --- | --- | --- | --- |
|  |  |  |  | p | p | p |
| Leukocyte | 1.23±2.05 | 1.69±2.77 | 2.80±0.62 | 0.109 | 0.285 | 0.285 |
| Red blood cell | 4.75±0.75 | 4.77±0.77 | 4.23±0.50 | 0.593 | 0.285 | 0.109 |
| Hemoglobin | 15.20±0.45 | 15.56±0.28 | 15.16±0.83 | 0.109 | 0.655 | 0.414 |
| Hematocrit | 41.53±4.84 | 43.76±6.35 | 35.00±7.07 | 0.285 | 0.285 | 0.109 |
| MCV | 87.80±6.82 | 92.30±12.05 | 82.53±12.36 | 0.285 | 0.285 | 0.109 |
| MCH | 32.53±5.40 | 33.20±5.23 | 36.30±6.15 | 0.102 | 0.109 | 0.109 |
| MCHC | 37.00±5.30 | 36.16±6.33 | 44.70±10.63 | 0.593 | 0.109 | 0.109 |
| Platelet | 265.66±51.03 | 294.66±70.51 | 289.66±106.95 | 0.109 | 0.593 | 1.000 |
| RDW-SD | 79.73±5.01 | 76.80±3.35 | 87.96±4.79 | 0.655 | 0.109 | 0.109 |
| RDW-CV | 25.86±0.90 | 23.96±3.82 | 30.30±3.43 | 0.109 | 0.109 | 0.109 |
| HbA1c | 5.40±0.36 | 5.50±0.51 | 5.33±0.49 | 0.414 | 0.414 | 0.102 |

**Table 2:** Astronauts Time-Dependent Correlation Analysis (correlation analysis)

|  | Astronaut A | Astronaut B | Astronaut C |
| --- | --- | --- | --- |
|  | p | p | p |
| Leukocyte | 0,3252 | 0,8078 | 0,3248 |
| Red blood cell | 0,3508 | 0,5456 | 0,2913 |
| Hemoglobin | 0,6667 | 0,0732 | >0,9999 |
| Hematocrit | 0,4246 | 0,8965 | 0,3036 |
| MCV | 0,5292 | 0,9341 | 0,3223 |
| MCH | 0,1919 | 0,1789 | 0,2531 |
| MCHC | 0,3401 | 0,9406 | 0,9421 |
| Platelet | 0,4605 | 0,1603 | 0,9421 |
| RDW-SD | 0,2086 | 0,7034 | 0,3333 |
| RDW-CV | 0,3534 | 0,8675 | 0,3419 |
| HbA1c | 0,7877 | >0,9999 | 0,6667 |

**Fig. 1.**
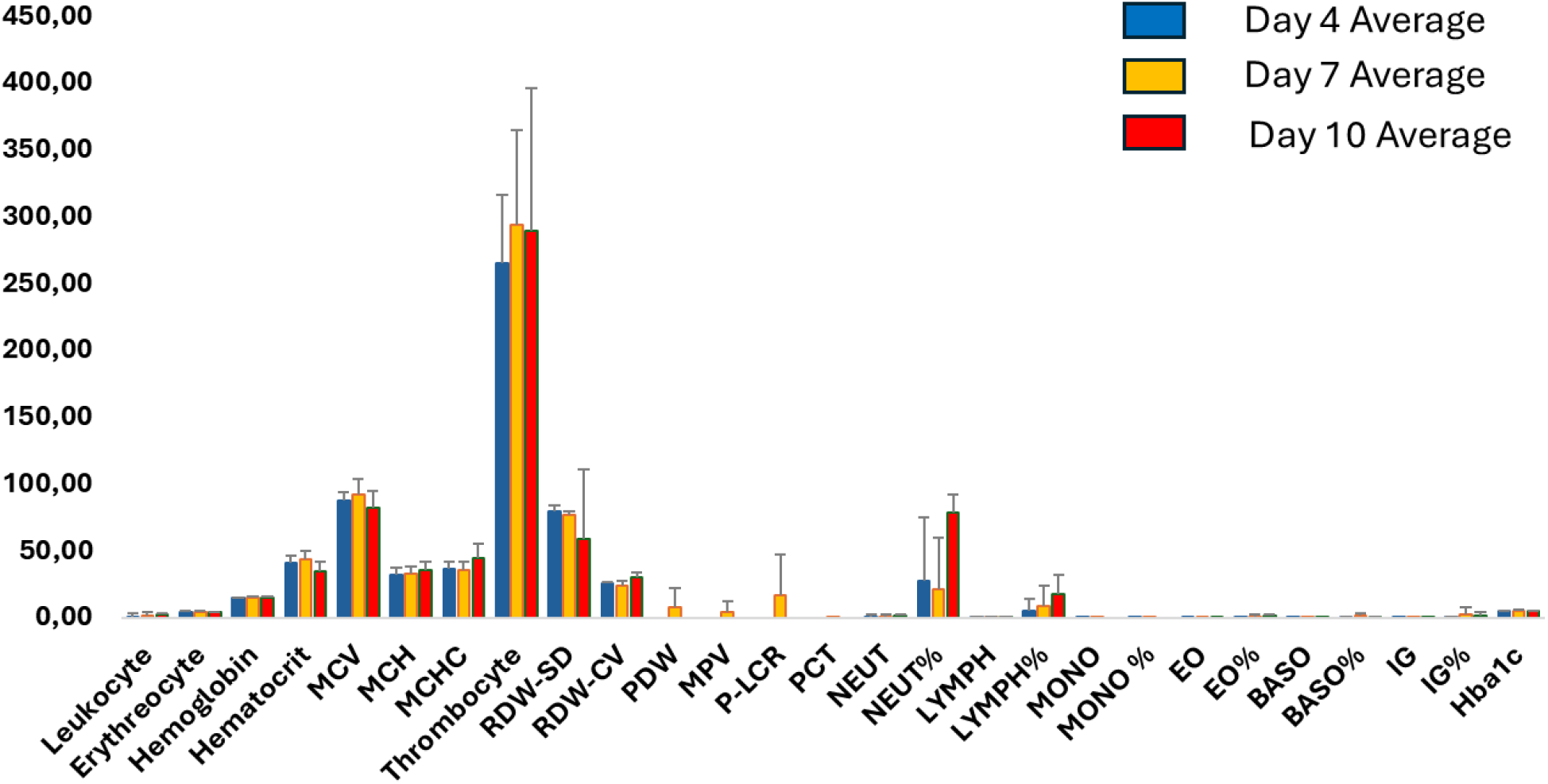
Hemogram Analysis Results of astronauts whose blood samples withdrawn in space in Day 4, 7, and 10.

Based on the time-dependent correlation analyses conducted for each astronaut, no statistically significant changes (**p > 0.05**) were observed in the measured parameters between **Day 4 and Day 10**. This finding suggests that the assessed biological processes did not exhibit substantial variation within this timeframe and that the evaluated parameters remained stable in the microgravity environment over the short term.

This study aimed to investigate potential changes in various haematological parameters by analysing blood samples collected from three different astronauts on days 4, 7, and 10 of their respective missions. The parameters analysed included leukocyte, erythrocyte, haemoglobin, haematocrit, MCV, MCH, MCHC, thrombocyte, RDW-SD, RDW-CV, PDW, MPV, P-LCR, PCT, NEUT, NEUT%, LYMPH, LYMPH%, MONO, MONO%, EO, EO%, BASO, BASO%, IG, and HbA1c. The findings revealed no statistically significant changes in any of the analysed parameters, suggesting that haematological parameters remain stable during long-duration space missions and indicating a physiological adaptation to the space environment.

### Total Cell Viability and Count Analysis

Next, we wanted to analyse the cell viability in astronaut blood samples. The (see Fig. 2) shows live cell counts measured on Day 4, Day 7, and Day 10 for three astronauts (A, B, and C). The average live cell count increased from Day 4 to Day 7, reaching a peak, followed by a decrease on Day 10 across all groups. Astronaut A consistently maintained the highest number of live cells throughout all time points, whereas Astronaut C had the lowest counts, particularly on Day 10, where a sharp decline was observed. Total cell counts also followed a similar trend, with an initial increase from Day 4 to Day 7. Also, by Day 10, non-significant reduction was observed, particularly in Astronaut C, which had the lowest total cell count. Astronaut A maintained the highest count, while Astronaut B showed intermediate results. The average cell viability percentage remained relatively stable from Day 4 to Day 7 but showed a slight decrease by Day 10. Astronaut A showed the highest viability across all time points, maintaining above 60%, while Astronaut C had the lowest viability, particularly on Day 10, where it dropped below 50%. The data suggest that while live and total cell counts initially increased, possibly due to adaptation to experimental conditions, a decline in both live cell count and viability by Day 10 indicates cellular stress or reduced proliferative capacity over time, especially in Astronaut C. Astronaut A showed a relatively more stable profile across all metrics, suggesting greater resilience under the given microgravity conditions.

**Fig. 2.**
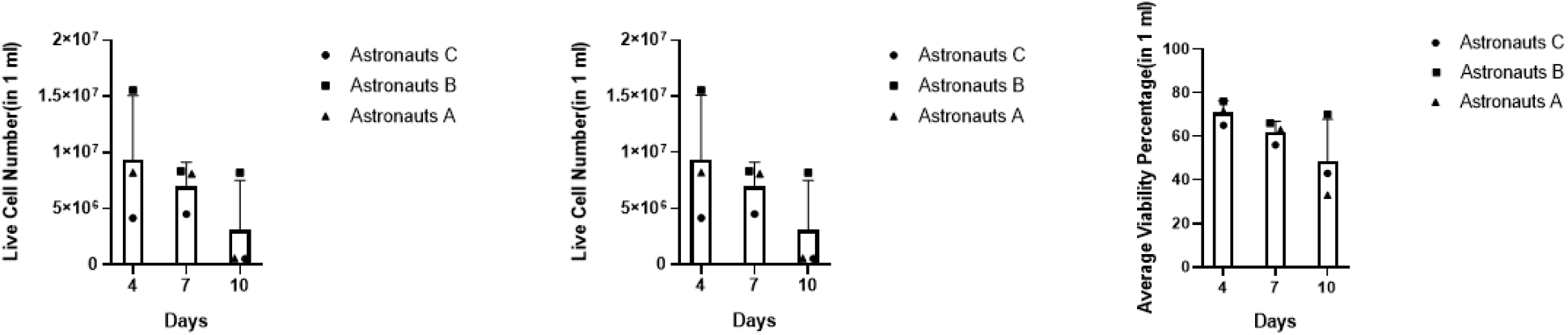
Viability, number of viable cells and total cells count graphs obtained because of cell count readings.

### T Cell Subpopulation Analysis

We compared various T cell subpopulations between astronauts and healthy donors at three time points: Day 4, Day 7, and Day 10. The subpopulations included in the analysis were TemEARLY (Early effector memory T cells), Tcm (Central memory T cells), Tn+Tscm (Naive and stem cell-like memory T cells), Temra (Effector memory T cells re-expressing CD45RA), and TemLATE (Late effector memory T cells) analysed by flow cytometry (see Fig.3.)

**Fig. 3.**
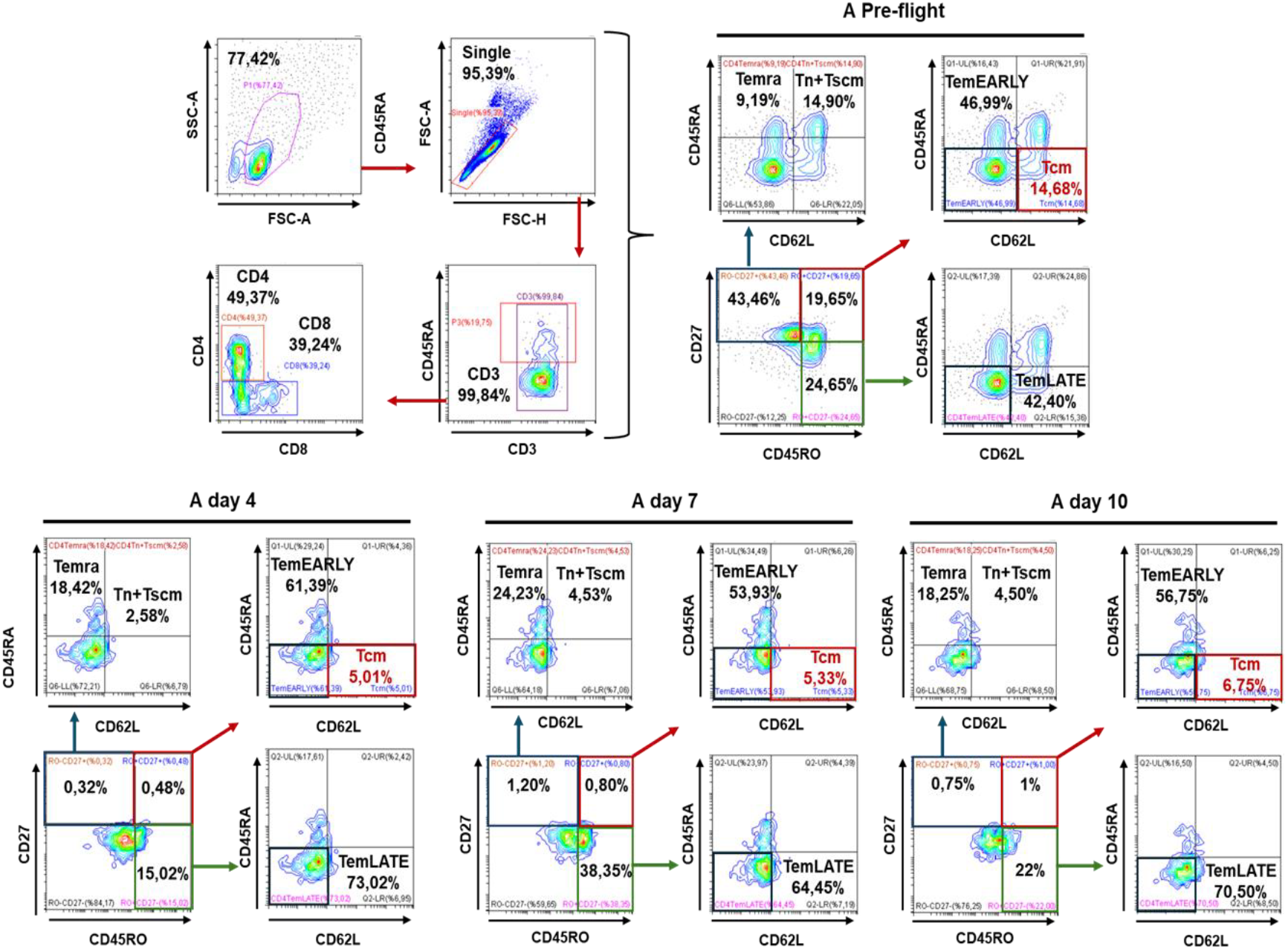
T cell subpopulations between astronauts and healthy donors at different time points (Pre-flight, Day 4, Day 7, and Day 10).

Next, we aimed to assess CD4+ T cell subpopulation analysis, including central memory T cells (TCM) and stem-cell like memory T cells (TSCM), measured at three time points: Day 4, Day 7, and Day 10 during spaceflight (see Fig. 4.). In day 4, the percentage of TemEARLY and TemLATE cells was high, with approximately 50% of the total CD4+ population comprising these effector memory subsets. Tcm and Tn+Tscm populations were minimal, constituting a small fraction of the total CD4+ population. Temra cells were present at low levels on Day 4. On the other hand, in day 7, TemEARLY cells slightly decreased in percentage, while TemLATE cells showed a steady increase, indicating a shift toward a more differentiated effector memory T cell population.

**Fig. 4.**
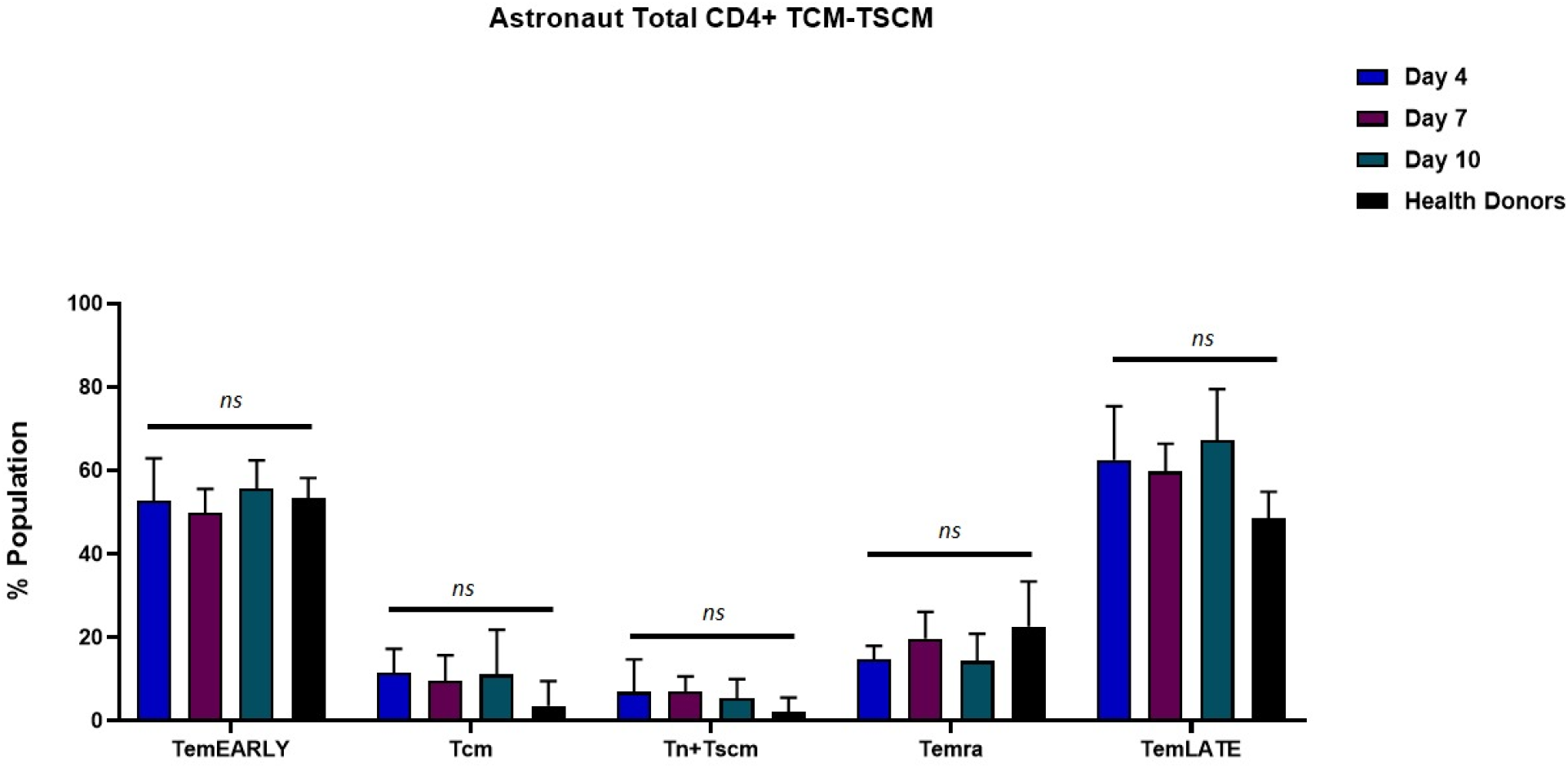
CD4+ T cell subpopulations, including central memory T cells (TCM) and stem-cell like memory T cells (TSCM), measured at three time points: Day 4, Day 7, and Day 10 during spaceflight

Tcm and Tn+Tscm populations remained low, but Temra cells exhibited a slight increase compared to Day 4. By Day 10, TemEARLY cells continued to decrease slightly, while TemLATE cells reached their highest levels, with over 60% of the population comprising late effector memory T cells. Tcm and Tn+Tscm populations remained low, like earlier time points, while Temra cells showed a notable increase, reflecting potential ongoing differentiation or immune activation as spaceflight progressed. The data suggest a shift from early effector memory T cells (TemEARLY) to late effector memory T cells (TemLATE) over the course of spaceflight. By Day 10, TemLATE cells constituted the majority of the CD4+ T cell population. Central memory T cells (Tcm) and stem-cell like memory T cells (Tn+Tscm) remained consistently low throughout all time points, suggesting that spaceflight conditions did not favour the expansion of these less differentiated memory subsets. The increase in Temra cells by Day 10 indicates potential ongoing immune activity or response to environmental stressors in space. These results reflect a shift in immune memory and differentiation dynamics during spaceflight, with a predominant focus on effector memory subsets, which could have implications for immune function during long-term space missions.

## 4. Discussion

The findings of this study highlight the substantial impact of microgravity on the immune system, particularly on T lymphocyte populations. Our observations of altered T cell subpopulations and cellular viability in astronauts during a 10-day space mission provide critical insights into how spaceflight can modulate immune function. Our data show substantial shifts in T cell subpopulations, blood cell profiles, and cell viability, reflecting the body’s adaptive responses to the unique stressors of spaceflight.

The hemogram analysis revealed key changes in blood cell populations during the mission. The most notable observation was the gradual increase in leukocytes (white blood cells) from Day 4 to Day 10, suggesting that microgravity may activate immune responses. Leucocytosis, observed by the increase in leukocyte counts on Day 10, might be due to the stress experienced in space, which can trigger immune activation. This increase in leukocyte count has been previously reported in studies that simulated spaceflight conditions, where the body responds to stress by activating innate immune pathways [1]. However, the erythrocyte count showed a slight decline by Day 10, which could indicate a reduction in red blood cell production or increased haemolysis in microgravity. This aligns with findings from other spaceflight studies that observed alterations in erythropoiesis and red cell mass during prolonged exposure to microgravity [6]. The non-significant reduction in haematocrit on Day 10 further supports the hypothesis of potential blood volume changes or fluid shifts in space. These alterations in haematological parameters, though subtle, highlight the need to monitor astronauts’ oxygen-carrying capacity and overall hematologic health during long-term missions [4]. The mean corpuscular volume (MCV) and mean corpuscular haemoglobin concentration (MCHC) remained relatively stable, though slight fluctuations in MCV from Day 4 to Day 10 may indicate cellular adjustments to the changing environmental conditions. Importantly, HbA1c levels, a marker of long-term glucose control, remained stable, suggesting that glucose metabolism was not significantly affected during the short duration of the mission. The analysis of cell viability and total cell counts provides additional insights into how immune cells adapt to microgravity. We observed an initial increase in live and total cell counts from Day 4 to Day 7, likely reflecting an initial proliferative response to microgravity. However, by Day 10, there was a notable decline in both live and total cell counts, particularly in astronaut C, suggesting that microgravity imposes cellular stress over time, ultimately reducing proliferative capacity and viability. This decline in cell viability aligns with previous studies that documented reduced cell proliferation and increased apoptosis in microgravity conditions [9]. The sharp decline in viability observed on Day 10 could be indicative of cellular exhaustion or the cumulative effects of stress, which has been reported in both simulated and real microgravity environments. The slight decrease in cell viability percentage from Day 7 to Day 10 further supports the notion that prolonged exposure to microgravity leads to cellular stress and reduced survival [2]. Interestingly, astronaut A maintained the highest live cell count and viability across all time points, suggesting potential individual variability in response to microgravity. This may be due to genetic factors, differences in immune resilience, or varying levels of adaptation to space conditions. Further research is needed to explore these individual differences and identify potential biomarkers of resilience to spaceflight-induced immune stress.

The non-significant elevation of effector memory T cells (TemEARLY and TemLATE) during spaceflight, as documented in our results, underscores a shift towards an activated immune state. This shift may be an adaptive response to the unique stressors encountered in the microgravity environment, including altered cellular signalling and reduced mechanical stress on cells. Previous studies have suggested that microgravity induces a hyper-responsive state in immune cells, which could predispose astronauts to inflammatory conditions or other immune-related health issues [3,6]. The persistent low levels of central memory T cells (Tcm) and stem cell-like memory T cells (Tn+Tscm) observed in this study may indicate a potential vulnerability in the immune system’s ability to respond to new antigens or fight off recurring infections. The reduction in these cell populations could lead to a compromised ability to form long-term immunological memory, which is essential for protecting against diseases over a lifetime [4]. The increase in late effector memory T cells (TemLATE) by Day 10 suggests a more differentiated state, potentially leading to an enhanced immediate defensive response but at the expense of proliferative capacity and longevity of the immune response. This observation aligns with findings from [1], who noted that spaceflight conditions could trigger early senescence of naive T cells, thereby accelerating the aging of the immune system.

The findings of this study indicate that, within the timeframe of the Axiom-3 mission, microgravity did not induce statistically significant changes in the haematological parameters and T cell subpopulations of the participating astronauts. The MESSAGE mission lasted only a limited number of days on the ISS, with blood samples collected on post-launch Day 4, Day 7, and Day 10. Previous studies suggest that immune dysregulation and haematological adaptations in microgravity may require prolonged exposure to space conditions. The duration of our study may not have been sufficient for significant physiological alterations to manifest, as previous research has shown more pronounced immune shifts during long-duration missions (>30 days). Astronauts undergo extensive pre-flight conditioning, including exercise regimens, nutritional optimization, and immune health monitoring, which could contribute to maintaining physiological stability in space. Additionally, onboard countermeasures such as exercise routines and controlled diets may have helped mitigate immune fluctuations and haematological shifts, preventing significant changes from occurring. While microgravity is known to influence immune function, other spaceflight-associated stressors such as radiation exposure, circadian rhythm disruption, and psychological stress may play a more significant role in immune changes over time. The absence of significant findings in this study suggests that short-term microgravity exposure alone might not be sufficient to induce acute immune shifts without the compounded effects of other stressors present in long-duration missions. With a cohort of only three astronauts, statistical power is inherently limited. The small sample size increases variability in individual responses and reduces the ability to detect significant trends. Future studies with a larger number of participants across multiple missions would provide a more comprehensive understanding of immune adaptations in space.

## 5. Conclusions

The implications of these findings are particularly relevant for long-duration space missions, where astronauts are exposed to microgravity for extended periods. Understanding the molecular mechanisms that drive these changes in T cell populations is crucial for developing strategies to mitigate adverse effects on astronaut health and maintain operational performance. Enhancements in spacecraft design, such as the inclusion of facilities to simulate gravity, may help in maintaining a more balanced immune response, as suggested by the simulated microgravity studies [9]. Although no statistically significant changes were detected in this study, further investigations are necessary to uncover potential molecular adaptations to microgravity. Next, transcriptome analysis will be performed to identify microgravity-associated genes, providing deeper insights into the regulatory pathways affected by spaceflight. Following gene identification, CRISPR-based knock-out T cells will be generated to determine the functional roles of these genes in immune modulation. This approach will allow for a more precise assessment of how microgravity influences immune function at the molecular level.

Understanding these gene functions is crucial for addressing the broader implications of spaceflight-induced immune changes, particularly in relation to pathogenicity, immune resilience, and cancer defence mechanisms. The ability to modulate immune responses in space could lead to novel therapeutic strategies for astronauts facing prolonged exposure to space environments. Additionally, insights from this study may have translational applications on Earth, contributing to new treatments for immunosuppression, infectious diseases, and cancer immunotherapy. Future research should expand on these findings by incorporating longer-duration missions, larger astronaut cohorts, and advanced molecular profiling techniques to develop targeted countermeasures for immune dysregulation in microgravity. Further research should focus on longitudinal studies involving astronauts to monitor the long-term effects of spaceflight on immune memory and resilience. Additionally, exploring countermeasures such as targeted therapies or interventions to support immune function could be pivotal. Developing on-ground analogues and model systems to simulate space conditions more accurately will also be essential in pre-mission training and post-mission recovery protocols [2].

## Acronyms/Abbreviations

MESSAGE: (Microgravity Associated Genetics) Science Mission
TemLATE: Transcriptional Enhancer-Like Activator
ISS: International Space Station
PBMC: Peripheral blood mononuclear cells
TCM: memory T cells
MCV: Mean corpuscular volume
MCHC: mean corpuscular haemoglobin concentration
TSCM: stem-cell like memory T cells
IL-2: interleukin-2
TNF-α: increased tumour necrosis factor-alpha
TRGENMER: Transgenic Cell Technologies and Epigenetic Application and Research Center

## Funding Acknowledgement

This study was supported by the Scientific and Technological Research Council of Türkiye (TÜBİTAK) and the Turkish Space Agency (TUA) through the Turkish Astronaut and Scientific Mission (TABM) Project (1007-KAMAG-121L002 / Agreement No: TABM-HZT-23-25. We would like to thank TÜBİTAK UZAY and TUA for their support and declare that none of the opinions and findings contained in this publication are the official views of TÜBİTAK and TUA.

## Acknowledgements

Manuscript presented at the International Astronautical Congress, IAC 2024, Milan, Italy, 14-18 October 2024.

